# Effects of genetic variability of CYP2D6 on neural substrates of sustained attention during on-task activity

**DOI:** 10.1101/831198

**Authors:** Roberto Viviani, Irene Messina, Julia E. Bosch, Lisa Dommes, Anna Paul, Katharina L. Schneider, Catharina Scholl, Julia C. Stingl

**Affiliations:** Institute of Psychology, University of Innsbruck, Austria; Psychiatry and Psychotherapy Clinic III, University of Ulm, Germany; Universitas Mercatorum, Rome, Italy; Research Division, Federal Institute for Drugs and Medical Devices, Bonn, Germany; Institute of Clinical Pharmacology, University Hospital of RWTH Aachen, Aachen, Germany

## Abstract

The polymorphic drug-metabolizing enzyme CYP2D6, which is responsible for the metabolism of most psychoactive compounds, is expressed not only in the liver, but also in the brain. The effects of its marked genetic polymorphism on the individual capacity to metabolize drugs are well known, but its role in metabolism of neural substrates affecting behavior personality or cognition, suggested by its CNS expression, is a long-standing unresolved issue. To verify earlier findings suggesting a potential effect on attentional processes, we collected functional imaging data while N=415 participants performed a simple task in which the reward for correct responses varied. *CYP2D6* allelic variants predicting higher levels of enzymatic activity level were positively associated with cortical activity in occipito-parietal areas as well as in a right lateralized network known to be activated by spatial attentional tasks. Reward-related modulation of activity in cortical areas was more pronounced in poor metabolizers. In conjunction with effects on reaction times, our findings provide evidence for reduced cognitive efficiency in rapid metabolizers compared to poor metabolizers in on-task attentional processes manifested through differential recruitment of a specific neural substrate.

## Introduction

The drug-metabolizing cytochrome P450 (CYP) enzyme CYP2D6 is responsible for the metabolism of several drugs, including most psychoactive compounds (Kirchheiner et al. 2004; Stingl and Viviani 2015). As a few other enzymes of this class, CYP2D6 is also expressed in neuronal cells in the brain (Miksys et al. 2002; Dutheil et al. 2008; Ferguson and Tyndale 2011), suggesting a possible role in metabolizing endogenous substrates (for review, see Stingl et al. 2013). The *CYP2D6* gene is characterized by the occurrence of several variants that have extreme effects on the functionality of the enzyme, which ranges from complete lack of function (poor metabolizers, 7% in Caucasians) to manifold enhancement (ultrarapid metabolizers, 2-5%, Stingl et al. 2013). The phenotypes are coded by genetic alleles that show huge interethnic variability (Ingelman-Sundberg 2005; LLerena et al. 2014). The variant coding the *CYP2D6*4* allele, which is the most frequently found allele leading to the poor metabolizer phenotype in Caucasian and rarely occurs in African populations, is likely to be the outcome of the introgression of a Neanderthal genetic variant (Ingelman-Sundberg et al. 2014; Steffens et al. 2019).

Because of the marked genetic polymorphism of CYP2D6 in man and the evidence for its expression in the brain, a long-standing question concerns the existence of possible behavioral phenotypes of this genetic variability (Bertilsson et al. 1989). Previous neuroimaging studies had provided preliminary evidence for a modulation of brain activity in association with the *CPY2D6* genotype (Kirchheiner et al. 2011; Stingl et al. 2012). Because this modulation has been previously detected in Stingl et al. (2012) in two very diverse tasks and involved parts of the alertness network such as occipital areas, we hypothesized an effect on a basic sustained attentional mechanism, such as one active while performing prolonged tasks (Stingl et al. 2013). Furthermore, such an effect was consistent with data from previous behavioral studies about effects of this genotype on cognition, showing the prominent involvement of sustained attention capacity (Peñas-LLedó et al. 2009). In a taxonomy of attentional processes, sustained attention is associated with endogenous efforts to focus on continuous task execution (Parasumaran and Davies 1984), in contrast with orienting to external stimuli or unfocussed mind wandering. Individual differences in sustained attention capacity constitute a stable trait across adult life (Parasumaran 1976) and are inheritable (Polderman et al. 2007).

On a molecular level, the constitutive role of CYP2D6 in biotransformation of neuroactive substrates is still unknown. Among others, it has been suggested that CYP2D6 may affect pathways in the endocannabinoid, serotonergic and dopaminergic neurotransmission (Bromek et al. 2010; Ozdemir et al. 2006; Stingl et al. 2013). With respect of the sustained attention hypothesis, a possible effect of this polymorphism on dopamine function is of particular interest since reward is known to facilitate attentional performance (Della Libera and Chelazzi 2006, 2011; Serences 2008; Anderson et al. 2013). Dopamine is known to be active not only when the opportunity for reward is detected in the environment (Schultz 2015), but also when sustaining on-task activity directed at rewards and the reactivity to contingencies involved in reward-related tasks (Nicola 2010; Schultz 2016). For this reason (and irrespective of any hypothesis on the effect of CYP2D6 on dopamine metabolism), it is important to control for effects of reward when assessing sustained attention function (Maunsell 2004).

To verify the sustained attention hypothesis, in the present functional imaging study we investigated the effects of *CYP2D6* genotype on the neural substrates elicited by a continuous performance task in which the amount of reward obtained from correct responses varied. The task was designed to assess possible effects of *CYP2D6* genotype on both cognition and reward sensitivity (Viviani et al. 2019). Each trial consisted of two phases: the brief initial appearance of a cue informing participants about the reward level of the trial (high or low), followed by a sustained attention task block in which participants worked to collect rewards by pressing one of two buttons, depending on the side on which dots appeared in a rapid sequence (for details, see the Methods section below). Cues that are informative about impending reward levels naturally command attention, as shown also by the known activation of neural substrates of dopaminergic networks they elicit (Berns et al. 2001; Knutson et al. 2001). Hence, the sustained attention phase that follows allows assessing activation of neural substrates at the net of the orienting attentional response elicited by the cue. As shown in a previous study, the on-task second phase activated prevalently right-lateralized parietal-prefrontal networks (Viviani et al. 2019) known to be involved in sustained attentional tasks (Paus et al. 1997, 2000; Corbetta and Shulman 2002). Reward levels modulated activity in this network as well as in known subcortical substrates of dopamine function, such as the nucleus accumbens and the ventral tegmental area/substantia nigra (Haber and Knutson 2010). Here, we assessed the effects of CYP2D6 polymorphism by testing the interaction of genotype with the contrasts between the phases of the trials (to assess attentional effects), between reward levels, and their interaction. Furthermore, we expected that, in case of an effect in the attention contrast, this would accrue mainly from a modulation of the second phase of the trial, in which the role of on-task attention is most prominent.

## Methods

### Participants

The present study involved 441 healthy volunteer participants of European origin who gave written informed consent after being recruited via placards in local facilities. After careful screening for psychiatric disorders (Sheehan et al. 1998), participants were admitted to the study if no exclusion criteria were met (current alcohol/drug addiction, anorexia, or current affective psychological disorders, pregnancy/lactation, severe acute or chronic diseases, psychoactive or long-term medication, metal implants, large tattoos or tattoos near the head). In total, 26 participants were excluded from the final analysis because of incidental clinical findings or imaging artifacts, equipment failure, failure to administer or complete the task, and for excessive movements during the task (see Data analysis below), leaving 415 individuals in the final sample (235 females, age 23.4±3.8 years). CYP2D6 activity scores of the excluded individuals did not differ from those included in the final sample (logistic regression, z = −0.032, n.s.). This study was conducted in conformance with the guidelines of the Declaration of Helsinki and was approved by the Ethical Committee of the University of Bonn (Nr. 33/15) and the Ethical Committee of the University of Ulm (Nr 01/15).

### Genotyping

DNA was extracted from EDTA blood using MagNA Pure LC DNA Isolation Kit (Roche Diagnostics, Penzberg) following the manufacturer’s instruction. Genotypes for *CYP2D6* were determined using PCR amplification with real-time PCR probes: rs35742686 (2549delA), rs1065852 (100C>T), rs3892097 (1846G>A), rs5030655 (1707delT) were examined using Taqman® SNP Genotyping Assays (ABI Life Technologies, USA). Briefly, 1-10 ng template in 9 μl Nuclease-free water were added to 10.0 μL of 2× TaqMan® Universal PCR master mix (ABI/Life technologies, USA) and 1.0 μL of a 20× combined primers and probes mix (ABI/Life Technologies, USA). The cycle conditions were: 95 °C for 10 min, 95 °C for 15 s, and 60 °C for 1 min. The last two steps were repeated 40 times. Allelic discrimination was performed by endpoint analysis. Rs16947 (2850C>T) and rs28371725 (2988G>A), were determined using LightSNP Assays (TIBMolBiol, Berlin). For this, 5 μL DNA (10 ng/μL) were amplified using Lightcycler® FastStart DNA Master HybProbe Mix (Roche, Germany) and the respective LightSNiP Assay (TIBMolBiol, Berlin). The following PCR protocol was applied: 10 min at 95°C, followed by 45 cycles for amplification: 10 s at 95°C, 10 s at 60°C, 15 s at 72°C. Melting curve analysis was performed to distinguish between the different genotypes. Gene duplications and deletions were analyzed with long range PCR using primer sets as described in Sistonen et al. (2005). In brief, 50 ng −500 ng genomic DNA were amplified using Long Range PCR Kit (Qiagen, Germany) and duplication or deletion specific primers (Table 1) following the manufacturer’s instruction. The cycle conditions were: initial denaturation with 93°C for 3 min, followed by 35 cycles for amplification: 93°C for 15 s, 62 °C for 30 s, 68°C 4 min.

**Table 1:**
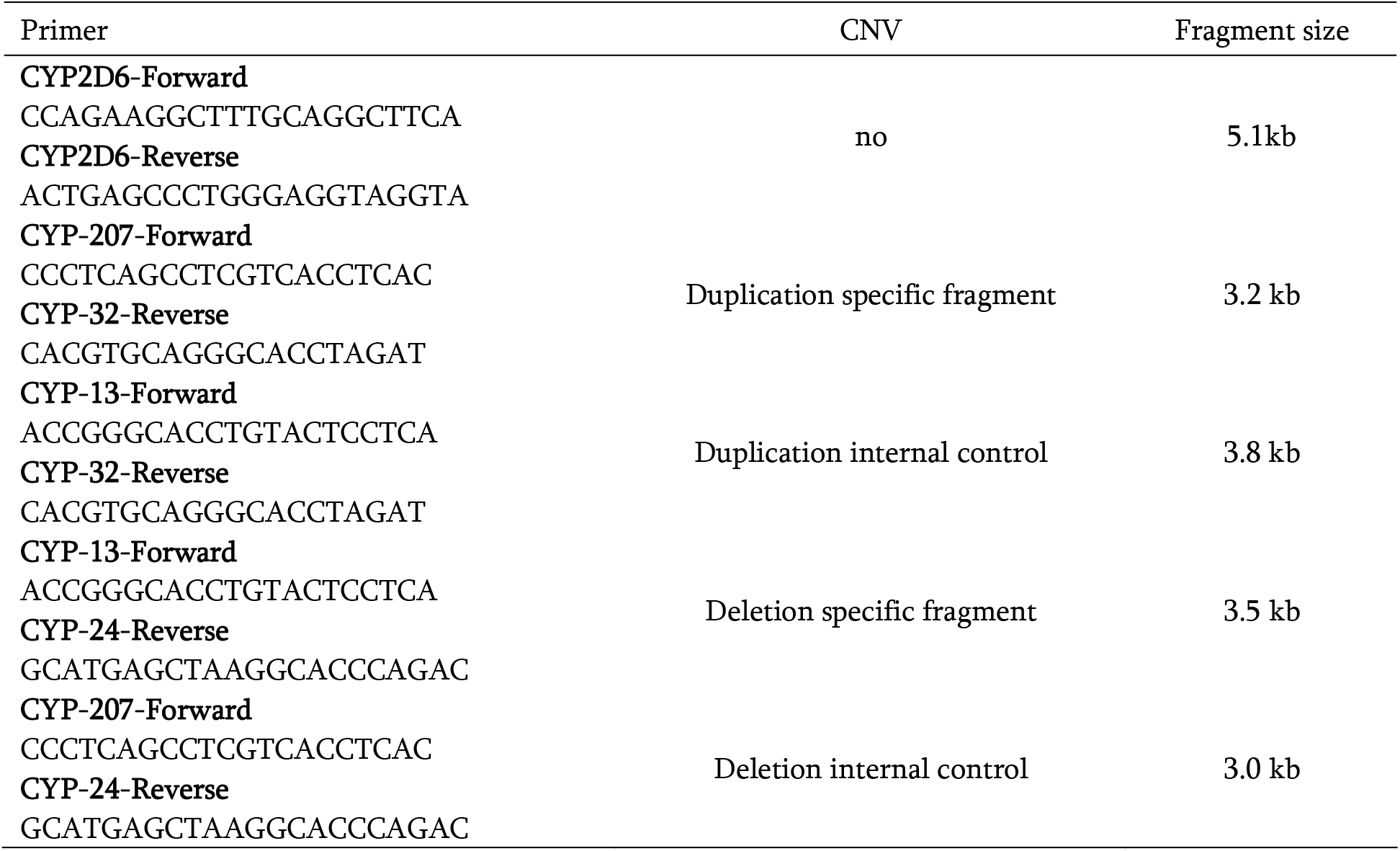
Primers used for determination of CYP2D6 CNV

PCR Products were separated on a 0.8% agarose gel and visualized after staining with DNA Stain Clear G (SERVA, Heidelberg) on a ChemiDoc imaging system (Biorad, Germany).

The genotyping results were used to derive the most likely haplotypes and to assign the most probable diplotype to each sample using the “CYP2D6 Allele Definition Table” provided by the Clinical Pharmacogenetics Implementation Consortium (CPIC) as reference (available online at https://www.pharmgkb.org/page/cyp2d6RefMaterials). To be as accurate as possible, the haplotype/diplotype analysis was performed manually by a team of physicians and pharmacists and additionally by a computerized algorithm. The haplotype/diplotype combination that was described best by the respective allelic constellation was chosen. In cases where more than one combination was consistent with the observed genotypes, the most prevalent haplotype/diplotype combination in the Caucasian population was assigned (e.g. *41 vs. *91 or *1xN/*2 vs. *1/*2xN). Based on the assigned diplotypes the CYP2D6 activity score was calculated as described by Gaedick et al. (2017).

### Functional imaging study task

Trials consisted of a cue announcing the level of reward for the trial, which could be either 1 or 20 cents at each correct response, and a rapid sequence of target dots appearing on the left or the right of the fixation point at fixed positions. Participants were required to press a left or a right button depending on the location the target dot (Figure 1). Each correct response always delivered either 1 or 20 cents, as announced by the preceding cues. The cue was displayed for 2 sec, was followed by a fixation cross for 3 sec, and by a block lasting about 15 sec during which 12 to 13 target dots were presented. Target dots appeared at irregular intervals according to an exponential schedule bounded between 800 and 1800 msec, and an average interval of about 1230 msec. The target dots remained on screen for a maximal duration of 800 msec, after which the award would not be collected. Trials were separated by pauses of 10 sec. during which a dimmed fixation cross was shown, for a total duration of 8 min for the whole task (16 trials). Cues and target dots belonging in different trial types were identifiable by color. Participants were conditioned on the cue color in a practice run prior to being positioned in the scanner. To match effort between the high and reward trials (e.g., ensure that participants would be working to collect the 1 cent rewards), the total reward would be paid only if participants collected a minimal sum of 20 Euro. Participants were told that they would need to work to collect the 1 cent rewards to reach that sum, and the sequence of trials was engineered so that the minimal sum would be reached towards the end of the experiment.

**Figure 1.**
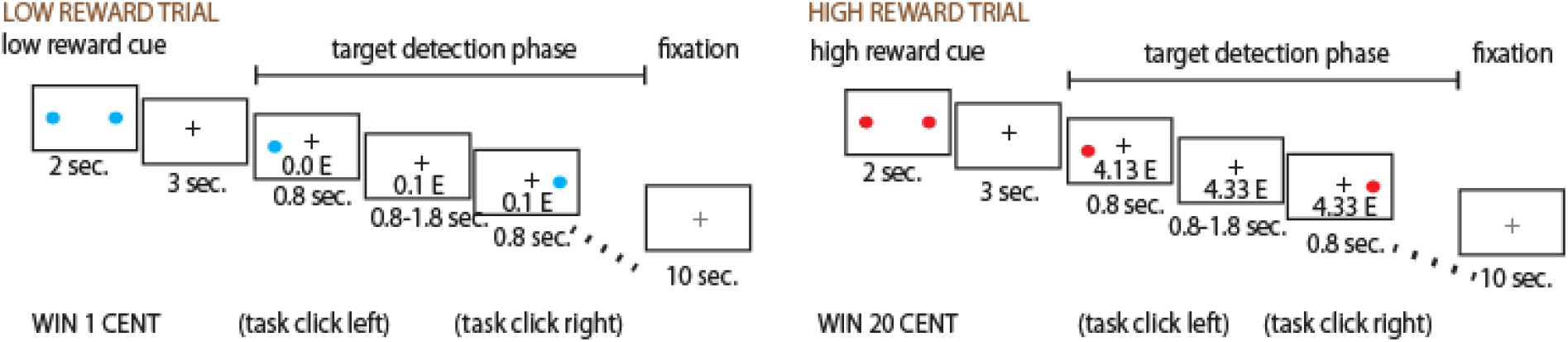
Schematic illustration of the task in the scanner. Participants had to click on the side on which a button appeared, here shown for two exemplary trials, one with low and one with high reward levels.

### Functional imaging data acquisition

Data were collected with a 3T Siemens Prisma (in the Psychiatry and Psychotherapy Clinic of the University, Ulm) and a 3T Siemens Skyra scanner (located on the premises of the German Center for Neurodegenerative Diseases, DZNE, Bonn), equipped with 64-channels head coils with a T2*-sensitive echo planar imaging sequence (TR/TE: 2460/30 msec), flip angle 82°, FOV 24 cm, 64×64 pixels of 3×3 mm in 39 2.5 mm transversal slices (in ascending acquisition order) with a gap of .5mm, giving an isotropic pixel size of 3mm. To adjust for shorter T2* in high susceptibility and iron-rich regions in the lower slices, echo time was gradually shortened by 8msec from slice 24 to slice 14, giving a TE of 22 msec in the first 14 slices acquired in the bottom of the volume, as described in Stöcker et al. (2006). Participants were individually screened for structural abnormalities in a T1-weighted structural image acquired during the session.

### Data analysis

Statistical analyses of behavioral data were conducted with the function lmer in the freely available packages lme4 (v1.0, Bates et al. 2015) in R (www.r-project.org/). Logistic and linear regression models (for hits and reaction times, respectively) included age, sex, site, reward levels, block and target number as confounding covariates, and the random effect of subjects to account for repeated measurements.

Neuroimaging data were analysed with the software SPM12 (Friston et al. 1995, Wellcome Trust Centre for Neuroimaging, http://www.fil.ion.ucl.ac.uk/spm/) running on MATLAB (The MathWorks). Data were first realigned to the first image to correct for head motion, normalized into the standard Montreal Neurological Institute (MNI) space and resampled to an isotropic voxel size 2 mm, and smoothed with a Gaussian kernel (8 mm FWHM). Data were regressed at the first level on a box-car function convolved with a canonical hemodynamic response function of fixed amplitude and on the realignment parameters as confounding covariates. Separate regressors were used to model the cue and the foraging phases in the low and high reward levels. A first-order autoregressive model accounted for the temporal autocorrelation of residuals at the first level. Estimates of contrasts of interest (detection of target dots vs. cues for on-task attention, high vs. low reward levels, the interaction, and the simple effects for these contrasts) were computed at the first level and brought to the second level to account for subjects as a random effect. At the second level, the model included CYP2D6 activity as estimated from the genotype, and age, sex of participants and site as confounding covariates. Note that both contrasts avoid the possible contribution of different baseline signal levels (Kirchheiner et al. 2011) because they compare different phases of the trial or different trials in the same run.

Reflecting the current transition between different approaches to correct for multiple testing, we based our inference on permutation techniques (Holmes et al. 1996, 8000 resamples). Cluster-level corrections were obtained by permutation for clusters defined at the uncorrected threshold *p* < 0.001. In the text, the size of cluster *k* is denoted in voxels (2×2×2 mm). Cluster threshold-independent tests were computed with the Threshold-Free Cluster Enhanced statistic (TFCE, Smith and Nichols 2009).

Overlays were prepared with the freely available software MRICroN (Chris Rorden, https://people.cas.sc.edu/rorden/mricron/install.html), while boxplots were drawn in MATLAB. Figure annotations were added with Adobe Illustrator.

## Results

### Genotyping

Results of the genotyping in the study sample (N = 415) are reported in Table 2. Activity scores did not significantly differ in age (*t*_414_ = −0.03) or sex (logistic regression, *z* = −0.5).

**Table 2.**
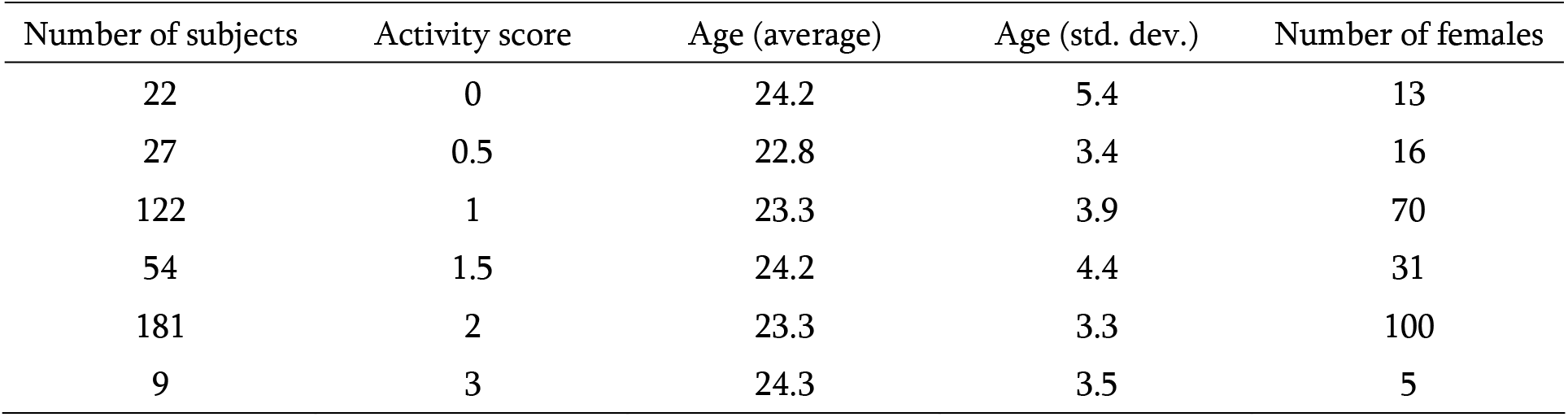
Results of genotyping

| Number of subjects | Activity score | Age (average) | Age (std. dev.) | Number of females |
| --- | --- | --- | --- | --- |
| 22 | 0 | 24.2 | 5.4 | 13 |
| 27 | 0.5 | 22.8 | 3.4 | 16 |
| 122 | 1 | 23.3 | 3.9 | 70 |
| 54 | 1.5 | 24.2 | 4.4 | 31 |
| 181 | 2 | 23.3 | 3.3 | 100 |
| 9 | 3 | 24.3 | 3.5 | 5 |

### Behavioral data

Participants had no difficulty in executing the task and gave the correct response in 99.4% of targets (range 86.6-100%). In the behavioral analysis, we verified that participants kept working at collecting rewards in both the low and high reward conditions, thus avoiding confounds arising from stopping to pay attention to the task and differences in motor activity. Reaction times averaged 423 msec (std. dev.: 60 msec) and were shorter in the high reward condition by about 4.5 msec. Although significant in this very large sample (*t*_408_ = 11.0, *p* < 0.001), this difference is very small when compared to the sluggishness of the BOLD response, which takes about 2 sec to rise (Buxton 2002).

There was no significant effect of *CYP2D6* genotype on reaction times (*t*_407_ = 1.0, n.s.). However, reaction times were affected by an interaction between *CYP2D6* genotype and reward levels (*t*_406_ = 3.2, *p* = 0.002, two-tailed). This interaction arose because low reward levels had a very small effect on reaction times in ultrarapid metabolizers (0.7 msec). The effect of low reward levels increased with decreasing CYP2D6 activity (2.4, 3.5., 3.2, and 5.3 msec for individuals with activity scores 2, 1.5, 1. and 0.5) reaching the highest effect sizes in poor metabolizers (7.5 msec). In summary, slowing of reaction times in association with increasing activity scores was not significant, but individuals with high activity scores were significantly less affected by reward levels. In term of frequency of correct responses, there was a trend-level reduced efficiency with increasing CYP2D6 activity scores (logistic regression, *z* = −1.72, p = 0.086, two-tailed).

### Neuroimaging data

In the analysis of the functional imaging data, we looked at the interaction of *CYP2D6* genotype with two main effects present in the task, attention type (sustained attention vs. orienting) and reward levels (high vs. low) and their interaction. In the factor attention type, *CYP2D6* genotype significantly modulated a fairly extensive cortical network comprising visual primary and secondary association areas in the occipital and parietal cortex (Figure 2 and Table 3), which extended medially into the posterior and middle cingulus. On the right hemisphere, activation in the inferior parietal cortex was accompanied by involvement of the precentral gyrus, extending towards the middle and inferior frontal gyri (Figure 2, rendered cortical surface on the right). Further prominent cortical activations were visible in the sensorimotor cortex and the insula. The boxplots shown in Figure 2 suggest that these effects were predominantly driven by the extreme metabolizer phenotypes. Subcortically, the posterior thalamus and the subthalamus were also modulated by *CYP2D6* genotype. In all these regions, the effect was positively associated with CYP2D6 activity as predicted by the genotype. There was no significant effect in the other direction. Further analyses revealed that this effect was due to a modulation of the fMRI signal in the sustained attention phase of the task, where activation was higher in individuals with high activity scores. In the sustained attention phase of the task, the effect of genotype was diffuse, but reached significance only in the superior parietal region (MNI coordinates x, y, z: −22 −62 58, *t* = 4.40, *p* = 0.042, peak-level corrected, and *p* = 0.140, TFCE correction). In contrast, no significant modulation of the signal was detected in the cue phase. However, we observed diffuse changes consistent with reduced activation in the cue phase with increasing activity scores.

**Table 3.** Interaction CYP2D6 activity and sustained attention (relative to orienting to cue)

| Cl. # | Location | MNI coord. | $k$ | $p$ clust.<br>(perm) | $p$<br>(TFCE) | $t$ | $p$ peak<br>(perm) |
| --- | --- | --- | --- | --- | --- | --- | --- |
| 1 | Lingual (BA 18) | -16 -64 -12 | 630 | 0.018 | 0.010 | 4.58 | 0.032 |
|  | Lingual (BA 18, 19) | -14 -76 -2 |  |  | 0.012 | 3.88 | 0.216 |
|  | Calcarine (BA 17) | -10 -84 6 |  |  | 0.014 | 3.62 | 0.362 |
| 2 | Lingual (BA 37) | -28 -48 -4 | 98 | 0.158 | 0.015 | 4.53 | 0.036 |
| 3 | Cuneus/Prec. (BA 7, 19) | 14 -70 34 | 762 | 0.013 | 0.009 | 4.52 | 0.039 |
|  | Precuneus (BA 7) | 4 -76 44 |  |  | 0.004 | 4.45 | 0.045 |
|  | Cuneus (BA 23) | -8 -76 50 |  |  | 0.011 | 3.88 | 0.216 |
| 4 | Insula | -42 8 4 | 196 | 0.080 | 0.019 | 4.41 | 0.049 |
| 5 | Sup. Parietal (BA 7) | 30 -54 64 | 147 | 0.109 | 0.013 | 4.36 | 0.059 |
| 6 | Sup. Orb. Front. (BA 11) | 32 62 -2 | 265 | 0.058 | 0.045 | 4.22 | 0.090 |
|  | Sup. Frontal (BA 10) | 30 60 16 |  |  | 0.048 | 4.08 | 0.135 |
| 7 | Mid. Orb. Front. (BA 11) | -28 40 -10 | 68 | 0.221 | 0.030 | 4.25 | 0.082 |
| 8 | Inf. Parietal (BA 40) | 50 -54 50 | 211 | 0.075 | 0.013 | 4.21 | 0.091 |
|  | Angular gyrus (BA 40) | 42 -52 38 |  |  | 0.014 | 3.77 | 0.237 |
| 9 | Precentral (BA 6) | -26 -20 46 | 125 | 0.130 | 0.015 | 4.04 | 0.149 |
|  | Precentral (BA 6) | -38 -16 56 |  |  | 0.019 | 3.34 | 0.555 |
| 10 | Mid. Cingulum (BA 23) | 10 -32 42 | 179 | 0.088 | 0.016 | 3.97 | 0.179 |
|  | Precuneus | 16 -44 44 |  |  | 0.018 | 3.64 | 0.348 |
|  | Mid. Cingulum (BA 23) | 10 -20 34 |  |  | 0.019 | 3.29 | 0.590 |
| 11 | Precentral (BA 6) | 40 -8 32 | 133 | 0.121 | 0.027 | 3.70 | 0.314 |
|  | Postcentral (BA 4) | 50 -6 32 |  |  | 0.028 | 3.64 | 0.345 |
| 12 | Postcentral (BA 3) | -60 -24 40 | 116 | 0.0140 | 0.019 | 3.47 | 0.463 |
|  | Supramarginal (BA 2) | -50 -22 50 |  |  | 0.019 | 3.41 | 0.505 |
| 13 | Ant. Hipp./Amy. | 24 -16 -12 | -- | -- | 0.035 | 2.69 | 0.913 |
| 14 | Subthalamus | -10 -22 -2 | 73 | 0.208 | 0.029 | 3.50 | 0.438 |
|  | Thalamus | -7 -22 5 |  |  | 0.034 | 2.74 | 0.895 |
|  | Ant. Hipp./Amy. | -20 -14 -12 |  |  | 0.029 | 3.37 | 0.532 |
| 15 | Subthalamus | 6 -12 -4 | 5 | 0.626 | 0.031 | 3.28 | 0.594 |
|  | Thalamus | 8 -19 1 |  |  | 0.033 | 2.80 | 0.871 |
| 16 | Nucl. Acc. | -14 6 -2 | 5 | 0.615 | 0.033 | 3.16 | 0.679 |
| 17 | Nucl. Acc. | 14 6 -2 | -- | -- | 0.035 | 2.93 | n.s. |
| 18 | Ventral Forebrain | -32 14 -12 | -- | -- | 0.034 | 2.86 | n.s. |
| 19 | Ventral Forebrain | 30 14 -8 | -- | -- | 0.037 | 2.93 | n.s. |
| 20 | Brainstem | 10 -24 -20 | -- | -- | 0.045 | 3.03 | n.s. |

Cl. #: cluster number; MNI coord: coordinates in Montreal Neurological Space; $k$ : cluster extent, in 2x2x2 mm voxels; $p$ clust. (perm), significance level, corrected at cluster level; $p$ (TFCE), significance level, TFCE correction; $p$ peak (perm), significance level, corrected at peak level with a permutation test. Inf., Mid., Sup., Ant.: inferior, middle, superior, anterior; Prec.: precuneus; Orb.: orbital; Front.: frontal; Hipp.: hippocampus; Amy.: amygdala; Nucl. Acc: nucleus accumbens.

**Figure 2.**
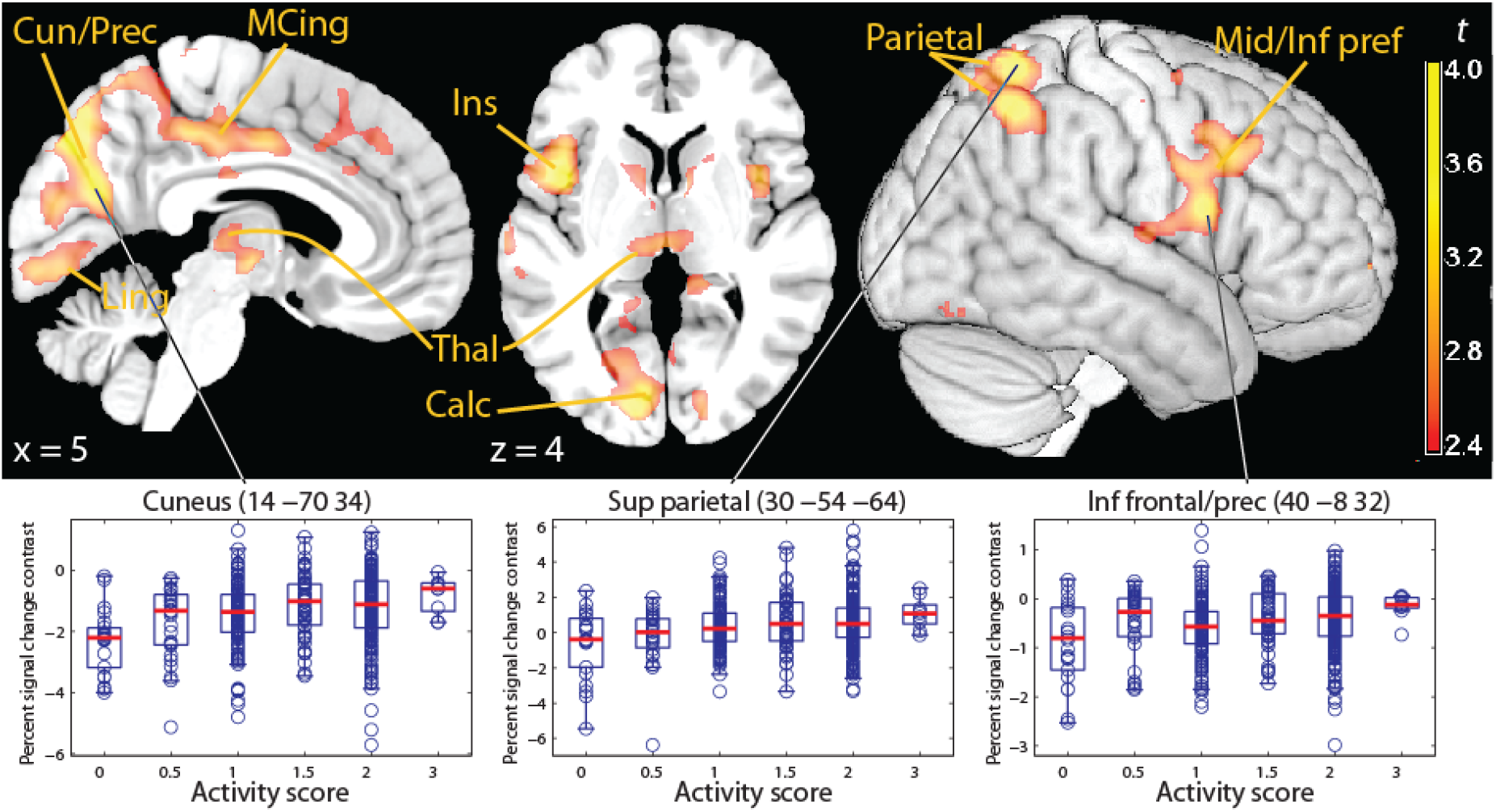
Cortical signal modulation by CYP2D6 activity (predicted from genotype) in the contrast sustained attention vs. orienting. Top row: parametric *t* map of the interaction CYP2D6 activity × attention type contrast, thresholded at TFCE significance (*p* < 0.05 corrected for the whole brain), overlaid on a template brain. The activation refers to higher activity in individual with high activity scores in the sustained attention phase of the task, relative to the cue phase, in the medial occipital areas (left) and in a right lateralized cortical network associated with spatial sustained attention tasks (right). Bottom row: boxplots of the signal contrast over CYP2D6 activity scores in selected voxels. Cun/Prec: cuneus/precunues; MCing: middle cingulus; Ling: lingual gyrus; Thal: thalamus; Ventr, Inf, Mid, Sup: ventral, inferior, middle, superior; frontal: frontal gyrus; prec: precentral.

In the factor reward levels, high activity scores were associated with reduced modulation of reward on brain activity in occipital and medial parietal cortical areas (Figure 3 and Table 4). This effect largely overlapped the previous contrast in the posterior occipital and parietal areas, but was less intense. This effect was due to individuals with higher activity scores showing higher activation in the low reward level trials (Fig. 3 right), thus limiting the extent of the modulation induced by reward levels.

**Table 4.** Interaction CYP2D6 activity and reward levels

| Cl. # | Location | MNI coord. | $k$ | $p$ clust. (perm) | $p$ (TFCE) | $t$ | $p$ peak (perm) |
| --- | --- | --- | --- | --- | --- | --- | --- |
| 1 | Inf. Occ. (BA 18) | -24 -90 -8 | 209 | 0.075 | 0.030 | -4.11 | 0.132 |
|  | Lingual (BA 18, 19) | -32 -84 -18 |  |  | 0.032 | -3.54 | 0.465 |
|  | Calcarine (BA 18) | -10 -96 -6 |  |  | 0.036 | -3.49 | 0.503 |
| 2 | Fusi. (BA37) | -38 -46 -24 | 66 | 0.233 | 0.045 | -3.72 | 0.337 |
| 3 | Inf./Mid. Occ. (BA 37) | -42 -70 -2 | 77 | 0.207 | 0.037 | -3.55 | 0.457 |
| 4 | Lingual (BA17) | -6 -80 -14 | 66 | 0.233 | 0.041 | -3.53 | 0.469 |
| 5 | Inf. Occ./Fusi. (BA 19) | 46 -66 -16 | 195 | 0.082 | 0.037 | -3.88 | 0.235 |
| 6 | Precuneus (BA 23) | -18 -58 32 | 116 | 0.142 | 0.033 | -3.96 | 0.194 |
| 7 | Precuneus (BA 23) | 20 -54 36 | 220 | 0.071 | 0.031 | -3.95 | 0.198 |

Cl. #: cluster number; MNI coord: coordinates in Montreal Neurological Space; $k$ : cluster extent, in $2 \times 2 \times 2$ mm voxels; $p$ clust. (perm), significance level, corrected at cluster level using a permutation test; $p$ (TFCE), significance level, TFCE correction; $p$ peak (perm), significance level, corrected at peak level using a permutation test. Inf., Mid.: inferior, middle; Occ.: occipital; Fusi: fusiform; Front.: frontal.

**Figure 3.**
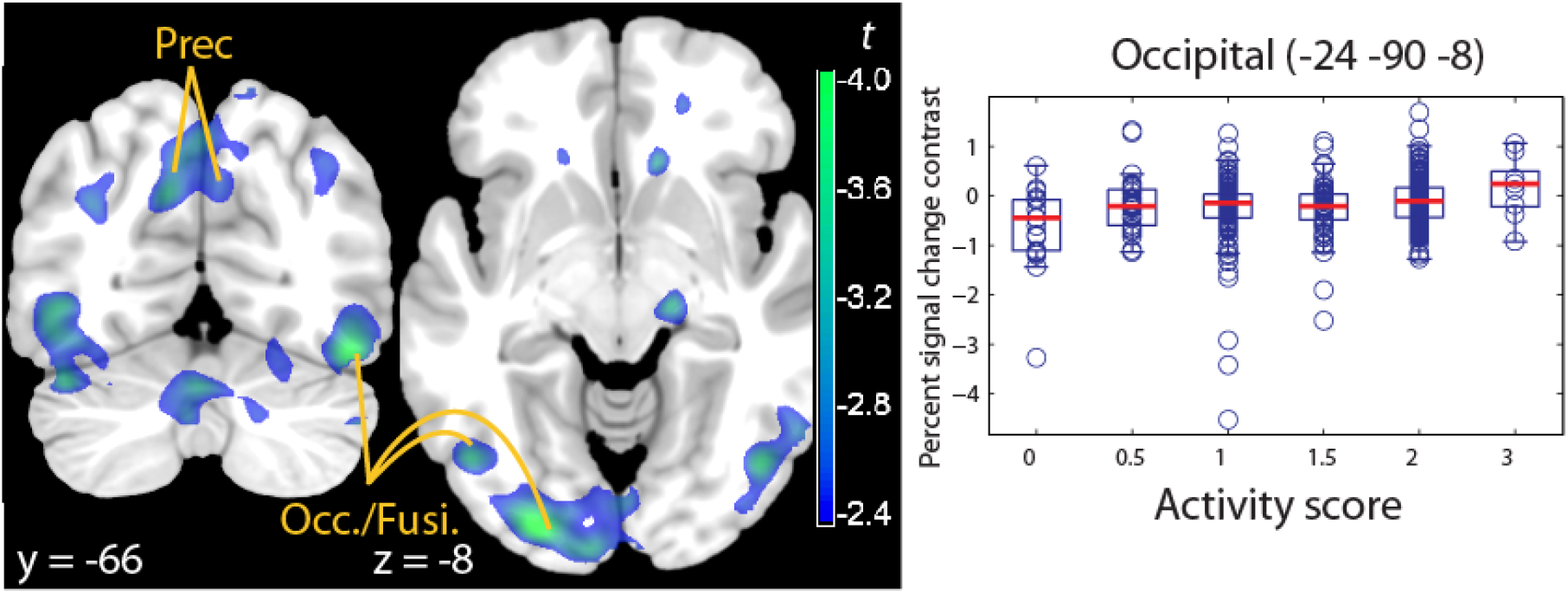
Cortical signal modulation by CYP2D6 activity (predicted by genotype) on the effect of reward. Left: Parametric *t* map of the interaction CYP2D6 activity × reward level contrast, thresholded at TFCE significance (*p* < 0.05 corrected for the whole brain), overlaid on a template brain. Higher activity in the high vs. ow reward contrast was negatively associated with CYP2D6 activity scores. Right: boxplots of the signal contrast for the low reward trials (relative to high), showing ultrarapid metabolizers (activity score 3) with no change in activity, while poor metabolizers (activity score 0) were affected by reward. The areas shown indicate less modulation by reward in occipital areas in individuals with high activity scores. Prec: precunues; Occ./Fusi.: inferior occipital and fusiform giri.

There was no significant interaction between genotype, attention type, and reward levels in the whole volume.

To better understand the role of individual reaction times on cortical activation patterns, we conducted a post-hoc analysis of the neuroimaging data and regressed them on the individual reaction times, taken as an index of efficiency of the individuals in performing the task (Honey et al. 2000; Binder et al. 2004). If the effects of *CYP2D6* genotype observed here are indicative of less task efficiency, we should observe changes in the same direction in individuals with long relative to short reaction times and in individuals with higher CYP2D6 activity scores. This is what we found (Figure 4). In the sustained attention relative to the cue phase contrast, high average reaction times were associated with higher activity diffusely across the cortex (similarly to the pattern seen for high CYP2D6 activity scores). This effect reached significance in the occipito-calcarine cortex (MNI coordinates x, y, z: 10, −86, 2, *t* = 3.88, *p* = 0.024, cluster-level corrected, and *p* = 0.061, TFCE correction: Figure 4 left, and −14, −80, 0, *t* = 4.19, *p* = 0.066, cluster-level corrected, and *p* = 0.060, TFCE correction). In the high vs. low reward contrast, high average reaction times were associated with a lower effect diffusely across the cortex (similarly to the pattern seen for high CYP2D6 activity scores). High reaction times produced lower differences in the signal between high and low reward levels, which reached trend-level significance in the same areas (12, −90, 4, *t* = −4.33, *p* = 0.080, cluster-level corrected, and *p* = 0.125, TFCE correction: Figure 4 right). In both cases, no significant effect was noted in the other direction of testing even at lenient thresholds.

**Figure 4.**
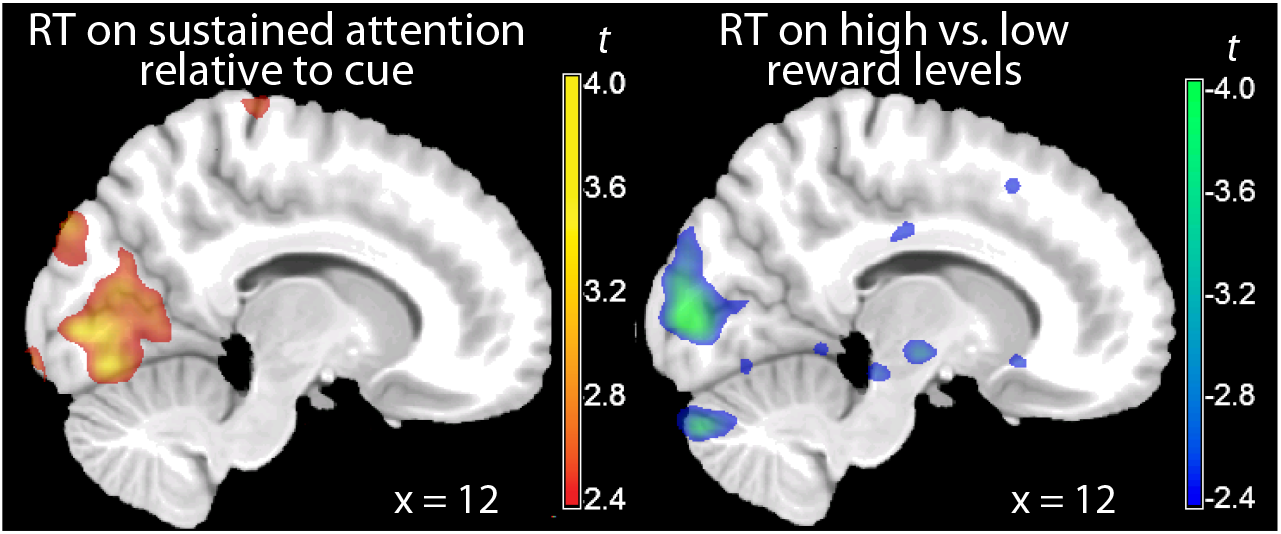
Effect of individual difference in reaction times on the cortical signal. Parametric *t* maps of the interaction between individual differences in reaction times (RT) and the fMRI contrasts attention type (left) and reward level (right).

## Discussion

The effect of the genetic polymorphism in CYP2D6 is usually found in patterns of metabolic ratios between drug substrates and metabolites built by this enzyme, reflecting its major functional role in the liver. In this study, in contrast, we focused on a possible role of this enzyme following from its expression in the brain. The present findings provide the strongest evidence to date for an effect of CYP2D6 activity on brain function in healthy humans. First, this is the first study in which the effect of *CYP2D6* genotype was tested on an *a priori* hypothesis about its effect on sustained attention with a task specifically designed for this purpose. Second, the effects observed in this task were large and inferentially valid independently of the method of testing. Third, the localization of the effects was consistent with the previous genetic neuroimaging studies of *CYP2D6*, revealing involvement of occipito-parietal cortical and subcortical areas during diverse tasks (Kirchheiner et al. 2011; Stingl et al. 2012). Fourth, the cortical changes were consistent with the behavioral effects on performance in a sustained attention task in a previous study (Peñas-LLedó et al. 2009).

In our findings, ultrarapid metabolizers activated more during the on-task phase of the trial, but were also less adaptive to the effect of reward levels, suggesting lower performance relative to poor metabolizers. This is consistent with the longer reaction times in the interaction with rewards levels observed in the behavioral data and provides new evidence on a cognitive trait associated with extreme metabolizer phenotypes. Previous behavioral data on smaller samples had described improved ‘energy levels’ (Bertilsson et al. 1989), less fatigability (Roberts et al. 2004), or larger sustained attention capacity in poor metabolizers (Peñas-LLedó et al. 2009). Poor metabolizers may be better at maintaining the focus of attention on a long task, as shown by the more efficient recruitment of spatial attentional networks in the present study.

The occipital areas, where the strongest effects of *CYP2D6* genotype were located in the attention contrasts, are known to be modulated by attentional effects (Corbetta et al. 1990; Gandhi et al. 1999). Other networks involved in the effect of genotype included specific cortical areas involved in sustained attention tasks, such as the middle and inferior right frontal gyri, as well as in the functionally relevant subcortical structures of the anterior striatum/basal forebrain (Paus et al. 1997, 2000). We also observed a selective involvement of the thalamus, consisting of a posterior portion (mainly localized to the lateral-posterior group and medial pulvinar) and the ventral-anterior portion (Najdenovska et al. 2018). A structural connectivity study by Behrens et al. (2003) has shown differential connectivity between thalamic nuclei and broad cortical areas. The portions of the thalamus associated here with CYP2D6 activity preferentially connect with the cortical areas detected in the same contrast, i.e. occipito-parietal and the middle and inferior prefrontal. The medial pulvinar, in particular, has been shown to be involved in visuo-spatial attention (Schmahmann 2003).

We also detected a weaker effect of *CYP2D6* genotype in the high vs. low reward contrast. Reward levels are known to affect activity in visual areas (Serences 2008), presumably because of facilitation of attentional efforts at high reward levels (Kurzban et al. 2013; Schultz 2016). The effect of *CYP2D6* genotype was due to higher activity in ultrarapid metabolizers in the low reward conditions, thus showing less modulation by reward. Although indicating that the attentional effects in poor metabolizers were adaptive to reward levels, these findings argue against CYP2D6 exerting an effect primarily through modulation of reward function. The strongest effects on neural activity observed in the present study affected the sustained attention phase of the task irrespective of reward levels. The effect of *CYP2D6* genotype observed in the reward contrast is rather consistent with a generic enhancement of attentional processes in poor metabolizers, as previously argued in Stingl et al. (2013), magnified by the known facilitation of reward on attention. This conclusion throws new light on the original observations on the cognitive phenotype of CYP2D6 polymorphism (Bertilsson et al. 1989), consistently with the notion that its action throughout life may lead to dispositional and cognitive effects.

## Acknowledgments

This work was supported by a Neuron-ERANET grant (‘BrainCYP’, Grant number BMBF 01EW1402B) and by collaborative grants from the Federal Institute for Drugs and Medical Devices (BfArM, Bonn, Germany, Grants No. V-15981/68502/2014-2017 and V-17568/68502/2017-2020). The authors declare no conflict of interest.

